# Light-mediated circuit switching in the *Drosophila* neuronal clock network

**DOI:** 10.1101/515478

**Authors:** M Schlichting, P Weidner, M Diaz, P Menegazzi, E Dalla-Benetta, C Helfrich-Förster, M Rosbash

## Abstract

The circadian clock is a timekeeper but also helps adapt physiology to the outside world. This is because an essential feature of clocks is their ability to adjust (entrain) to the environment, with light being the most important signal. Whereas Cryptochrome-mediated entrainment is well understood in *Drosophila*, integration of light information via the visual system lacks a neuronal or molecular mechanism. Here we show that a single photoreceptor sub-type is essential for long day adaptation. These cells activate key circadian neurons, namely the lLNvs, which release the neuropeptide PDF. Using a cell-specific CRISPR/Cas9 assay, we show that PDF directly interacts with neurons important for evening (E) activity timing. Interestingly, this pathway is specific for light entrainment and appears to be dispensable in constant darkness (DD). The results therefore indicate that external cues cause a rearrangement of neuronal hierarchy, which is a novel form of plasticity.

## Introduction

Circadian clocks evolved as an adaptation to the continuous change of day and night and are believed to provide organisms a fitness advantage. The underlying molecular machinery includes a transcriptional-translational feedback loop, which generates oscillations of clock gene expression with an endogenous period close to 24 hours (circa=about; dies=day) (Hardin, 2011). This period is approximately 24.2h in humans, whereas a *Drosophila* period was reported to be 23.8h (Czeisler et al., 1999; Dubowy and Sehgal, 2017). A key feature of circadian clocks is the ability to entrain to the 24h environment. This means that the human clock has to be accelerated by about 0.2h each day, whereas this *Drosophila* clock has to be slowed down to the same extent. To do so, clocks must integrate external cues, so called zeitgebers, which are used to synchronize the molecular and physiological properties of the organism (Golombek and Rosenstein, 2010).

The most important zeitgeber is light. In mammals, a combination of the traditional photoreception pathway (rods and cones) and the circadian photoreceptor melanopsin in retinal ganglion cells allows for fine-tuning of clock synchronization (Berson et al., 2002; Hattar et al., 2002; Lucas et al., 2012). Similarly, *Drosophila* uses the visual system and the circadian photoreceptor Cryptochrome (CRY) for light synchronization (Rieger et al., 2003; Stanewsky et al., 1998). CRY-mediated entrainment is well understood in *Drosophila*, whereas less is known about the mechanism of entrainment via the visual system. It consists of seven eye structures: three ocelli, two Hofbauer-Buchner-eyelets and two compound eyes (Hofbauer and Buchner, 1989).

The compound eye consists of approximately 800 ommatidia, each harboring 8 photoreceptor cells (Rs): R1-6 are located in the periphery and span the whole depth of the ommatidium. These cells were previously shown to be important for motion vision and express Rhodopsin 1 (Rh1) (Yamaguchi et al., 2008). In the center, R7 is located above R8. These cells have a complex expression pattern of Rh4 and/or Rh3 in R7 and Rh5 or Rh6 in R8 (Rister et al., 2013). How light information is conveyed from these photoreceptor cells to the circadian clock is not well understood.

The *Drosophila* clock neuron network consists of 150 clock neurons distributed in the lateral and dorsal parts of the brain. Recent electrophysiological results suggest that the visual system is able to activate an array of circadian clock neurons (Li et al., 2018), e.g., it can activate the small ventral lateral neurons (sLNvs), an important center for morning (M) activity (Grima et al., 2004; Stoleru et al., 2004). Furthermore, the visual system increases neuronal firing in the large LNvs (lLNvs), the arousal center within the circadian network (Shang et al., 2008). The 5^th^ sLNv and the NPF+ LNds, previously implicated as necessary for evening (E) activity (Hermann et al., 2012; Rieger et al., 2006, 2009), also increase their firing rates in response to visual system stimulation (Li et al., 2018). In addition, the visual system activates several dorsal neurons (DN), which were recently implicated in connecting the circadian clock to central brain sleep centers (Guo et al., 2018; Lamaze et al., 2018). These data suggest that visual input is integrated into the clock network in a parallel fashion, which contradicts a master-oscillator point of view (Li et al., 2018). The latter posits that these are the pigment-dispersing factor (PDF)-expressing neurons (sLNvs and lLNvs), which receive light input and release PDF upon illumination, thereby adjusting their downstream target neurons to the LD cycle (Yoshii et al., 2016).

To investigate the impact of the visual input pathway at the behavioral and neuronal level, we investigated fly behavior under long day conditions. Long days cause plastic changes in fly behavior: In standard light-dark cycles of 12h light and 12h darkness (LD 12:12), flies show a bimodal activity pattern with a M anticipation peak around lights-on and an E anticipation peak around lights-off; this results in a phase relationship of approximately 12h between the two peaks. Flies are able to adjust to longer photoperiods by delaying the E peak, which reduces the potentially harmful impact of hot summer days (Rieger et al., 2003). Previous experiments implicated the visual system in long day adaptation: Flies lacking the compound eyes fail to appropriately adjust their peak timings (Rieger et al., 2003). It is still unknown however which receptors and which neuronal pathways are involved in this adjustment.

Here we show that R8 of the compound eyes is essential for long day adaptation. These photoreceptor cells connect to the PDF-containing lLNvs and trigger the release of this neuropeptide. Using a cell-specific CRISPR/Cas9 strategy, we demonstrate that light-mediated PDF directly signals to the PDF receptor (PDFR) on E cells and hence delays E activity. The data implicate a mammal-like structure of clock entrainment, with the visual system activating PDF-expressing clock neurons. Our data further indicate a prominent shift of PDF targets between LD and DD conditions as well as a more quantitative reorganization of neuronal dominance within the clock network by changes in photoperiod.

## Material and Methods

### Fly strains and rearing

The following fly lines were used in this study: CantonS, *w^1118^*, *cli^eya^ (Bonini et al., 1993)*, *ninaE^5^* (BL 3545), *rh3^1^rh4^1^* (Vasiliauskas et al., 2011), *rh5^2^;rh6^1^* (Yamaguchi et al., 2008), *rh5^2^;rh3^1^rh4^1^rh6^1^* (Schlichting et al., 2014), *rh6-GAL4* (Sprecher and Desplan, 2008), *rh5-GAL4* (Mazzoni et al., 2008), *rh3-GAL4* (Wernet et al., 2006), *rh4-GAL4* (Wernet et al., 2006), UAS-*Kir2.1* (Baines et al., 2001), UAS-*HID* (BL 65403), *pdf(M)-GAL4* (Renn et al., 1999), *pdf^01^* (Renn et al., 1999), UAS-*pdf*-RNAi (BL 25802), UAS-*pdf* (Renn et al., 1999), UAS-*dcr2* (BL 24646), *R6-GAL4* (Helfrich-Förster et al., 2007), *c929-GAL4* (Hewes et al., 2003), UAS-*DenMark* UAS-*syt.eGFP* (BL 33065), *clk856-GAL4* (Gummadova et al., 2009), *mai179-GAL4* (Grima et al., 2004), *Spl-E-cell-GAL4* (Guo et al., 2017), *han^5304^*;UAS-*pdfr* (Hyun et al., 2005; Mertens et al., 2005), UAS-*Cas9.P2* (BL 58986), *w;CyO/Sco;MKRS/TM6B* (BL 3703). All flies were raised on standard cornmeal medium in LD12:12 at 25 degrees.

### Behavior recording and analysis

Single 2-6 days old male flies were transferred into glass tubes with food (1% agar and 4% sucrose) on one end and a plug to close the tube on the other end. The glass tubes were placed in *Drosophila* activity monitors (DAM) and a computer measured the number of light-beam interruptions caused by the fly in one minute intervals. Behavior was either recorded in light boxes within a climate controlled chamber or within incubators at constant temperature.

All flies were entrained for one week at LD 12:12. For photoreceptor mutants, flies were exposed to either LD 14:10 or LD 16:8 in week 2 and either to LD 18:6 or LD 20:4 in week 3. To investigate clock neuron interactions, flies were subjected to LD 20:4 after entrainment in LD 12:12 for one week. To determine free-running behavior, we entrained flies in LD 12:12 for at least 5 days and transferred them into constant darkness (DD) for at least 7 days.

Behavior analysis was performed as described in (Schlichting and Helfrich-Förster, 2015). We first generated actograms using ActogramJ (Schmid et al., 2011) and calculated average activity profiles out of the last 4 days of each light condition to allow for proper entrainment. We then generated single-fly average days and determined the timing of M and E peaks manually. Boxplots of single-fly values were generated to show the timing and the distribution of the data. Free-running periods were determined using chi^2^ analysis. Statistical analysis was performed using two-way ANOVA (StataSE 15), one-way ANOVA followed by post-hoc Tukey comparison (astata) or student’s t-test (Excel).

### Fly line generation

We generated a UAS-*PDFRg* fly line following the protocol of (Port and Bullock, 2016). In short, we digested the vector pCFD6 (addgene #73915) with BbsI. We PCR amplified two fragments carrying three independent guides for PDFR using Q5-polymerase (New England BioLabs, NEB) and performed a Gibson assembly (NEB). Positive clones were sent for injection to Rainbow Transgenic Flies (Camarillo, CA, USA) and inserted into the attP1 landing site on the second chromosome (BL 8621). W+ flies were balanced using BL3703 and kept as stable stocks above CyO. The following guides were used:

Guide 1: TCGAACATTCTCGACTGCGG
Guide 2: TGCTGGCCACCCACTCCGGC
Guide 3: CCTACATAGACATTGCCAGG

### Immunohistochemistry

#### Brain staining

2-6 days old male flies were fixed for 2h 45 min in 4% paraformaldehyde (PFA) in phosphate-buffered saline including 0.5% Triton X (PBST) at room temperature (RT). After rinsing 5x 10 min each with PBST, we dissected the brains and blocked with 5% normal goat serum (NGS) in PBST. We applied primary antibodies overnight at RT. The following antibodies were used: anti-PDF (1:1000, C7, Developmental Studies Hybridoma Bank (DSHB)), anti-GFP (1:1500, abcam, ab13970), anti-dsRed (1:1000, Living Colors DsRed Polyclonal Antibody, Takara). After rinsing 5x 10 min each in PBST, we applied secondary antibodies (Invitrogen, 1:200 dilution) for 3h at RT. After washing 5x 10 min each in PBST, brains were mounted on glass slides using Vectashield (VECTOR LABORATORIES INC., Burlingame, CA, USA) mounting medium.

#### Retina staining

2-6 days old male flies were fixed for 2h 30 min in 4% PFA in PBST at RT. After washing 5x 10 min each with PBST, retinas were dissected in PBST and blocked in 5% NGS in PBST for 1h at RT. Primary antibodies were applied for 2 nights at RT: anti-Rh1 (1:30, 4C5 DSHB), anti-Rh6 (1:2000, provided by C. Desplan, (Tahayato et al., 2003)), anti-chaoptin (1:50, 24B10 DSHB) and anti-GFP (1:1500, abcam, ab13970). After rinsing 5x 10 min each in PBST, we applied secondary antibodies (Invitrogen, 1:200 dilution) for 3h at RT. After washing 5x 10min each in PBST, retinas were mounted on glass slides using Vectashield (VECTOR LABORATORIES INC., Burlingame, CA, USA) mounting medium.

#### Imaging

All images were obtained using either a Leica SPE or a Leica SP5 confocal microscope. All brains/retinas were scanned using sections of 2 um thickness. Contrast and brightness were adjusted using FIJI and Photoshop CS5 extended.

### Expansion Microscopy

For expansion microscopy we applied the Pro-ExM protocol described in (Chen et al., 2015) as modified in (Guo et al., 2018). In short, after applying the regular IHC protocol described above, brains were transferred into the anchoring solution (Acryloyl X-SE, Life technologies A20770, 1:100 in PBS) for 24h. We rinsed the brains 3x 10 min each in PBS, before the gelling solution was added. We incubated the brains for 45 min on ice before transferring them into the gel chamber, in which they were incubated at 37°C for 2h. After the gel solidified, we trimmed away the excess gel material and applied the digestion buffer (ProteinaseK, 1:100 in PBS) for 24h. Subsequently, we washed the brians 3x 20 min each in ddH_2_O and placed the brains on a glass bottom culture dish (MatTek Corp, P35GC-0-14-C) with H_2_O. We generated the brain images using the Zeiss LSM 880 confocal microscope using z-stacks of 1 um. Image acquisition was performed using FIJI.

## Results

### The compound eyes are essential for long day adaptation

The locomotor activity of flies is controlled by their clock neuron network, which causes a bimodal pattern. In wild type (WT) flies (CantonS) under standard LD 12:12 conditions, the M peak of activity coincides with lights-on and the E peak with lights-off, respectively (Fig. 1A). In long photoperiods, the phase relationship between M and E peak increases showing plasticity in clock-controlled behavior (Fig. 1A and 1D) (Rieger et al., 2003). The M peak does not diverge from lights-on (Fig. 1B), whereas the E peak delays with increasing photoperiod (Fig. 1C), demonstrating that a delay of E activity is responsible for the enhanced phase relationship of the peaks (Fig. 1D). Notably, the E peak does not follow lights-off under all light conditions: Whereas it coincides with lights-off at LD 12:12 and LD 14:10, it occurs during the light phase at even longer photoperiods, resulting in a maximal phase relationship of 16.4 ± 0.3 h. Given that the E peak is the dominant factor for defining the phase relationship and the M peak is also less pronounced in some of the mutants (Fig. 1G), we focus on E peak timing as a surrogate for phase.

**Figure 1.**
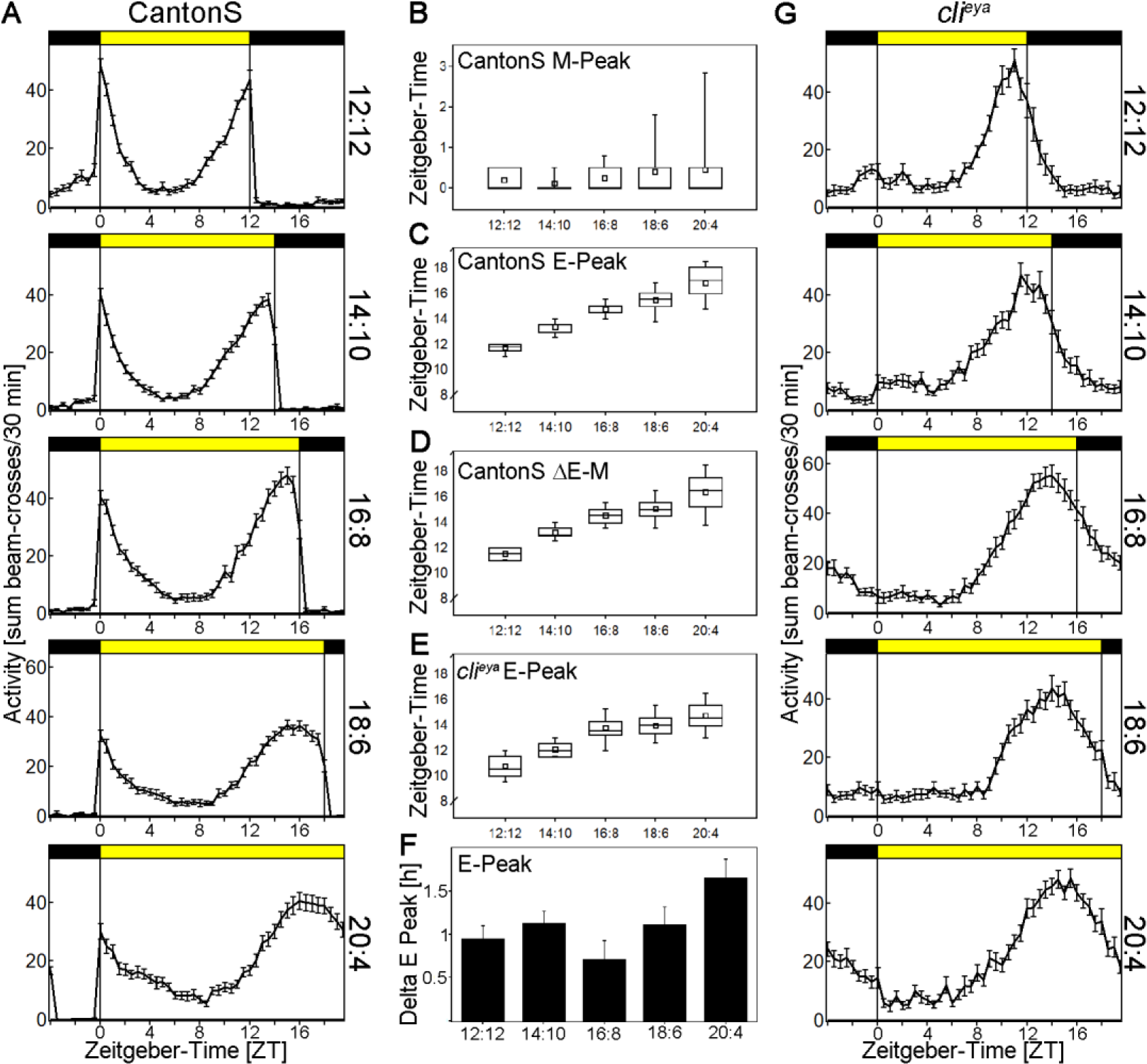
The compound eyes contribute to long photoperiod entrainment. **A** Average activity profiles of CantonS flies from LD 12:12 (top) to LD 20:4 (bottom). Daylight period increases by 2 hours per light condition. Flies show a bimodal activity pattern at every light condition with a broader E peak at longer photoperiods. **B** Timing of Morning (M) activity peak of CantonS flies at equinox and long day conditions. The M peak is tightly coupled to lights-on. It shows a tendency to delay with increasing day-length, which is not significant (F_(4,146)_=1.8064, p=0.1307). **C** Timing of Evening (E) activity peak of CantonS flies at equinox and long day conditions. Timing of the E peak delays with increasing day length, but does not follow lights off (F_(4,146)_=177.09, p<0.001). **D** Phase-relationship of M peak to E peak in CantonS at equinox and long day conditions. Due to the delay of the E peak timing, also the phase relationship increases with increasing daytime (F_(4,146)_=139.49, p<0.001). **E** Timing of E activity peak of *cli^eya^* flies at equinox and long day conditions. One-way ANOVA shows that *cli^eya^* flies delay their E peak timing with increasing daytime (F_(4,137)_=72.25, p<0.001). Two-way ANOVA shows a significant difference between CantonS and *cli^eya^* flies (F_(1,283)_=149.23, p<0.001) and a significant interaction between genotype and photoperiod, suggesting differential regulation of long day adaptation in the two genotypes (F_(1,283)_=3.85, p=0.0046). **F** Difference of E peak timing between CantonS and *cli^eya^* flies depending on day length. One-way ANOVA reveals a significant difference between the two genotypes, which is in agreement with the interaction of genotype and photoperiod described in E (F_(4,137)_=3.2766, p=0.0134). The biggest difference between the genotypes was found in LD 20:4. **G** Average activity profiles of *cli^eya^* flies from LD 12:12 (top) to LD 20:4 (bottom). Daylight period increases by 2 hours per light condition. *Cli^eya^* flies show a bimodal pattern in LD 12:12 which turns into unimodal behavior under long days with an early E peak timing.

To investigate the effect of the compound eyes on long day adaptation, we used *cli*^*eya*^ mutants lacking the compound eyes but retaining ocelli and Hofbauer-Buchner-eyelets (Schlichting et al., 2014). Even in LD12:12, the E peak is uncoupled from lights-off and is significantly advanced compared to WT flies (Fig. 1E and 1G). Even though eyeless flies adjust their E peak to long photoperiods (Fig. 1E and 1G), they fail to delay like WT flies, resulting in an approximately 1.5h maximally advanced E peak timing under LD20:4. We calculated ∆E-peak between CantonS and *cli*^*eya*^ mutants and found that this difference also depends on photoperiod: The longer the photoperiod, the bigger the difference between CantonS and *cli*^*eya*^, resulting in a maximal difference in LD20:4 (Fig. 1F). Therefore, we use this extreme photoperiod in the rest of this study to further investigate eye-mediated long day adaptation.

### Receptor cell 8 is responsible for *eyes absent* phenotype

The compound eyes are comprised of approximately 800 ommatidia. Each ommatidium contains 8 photoreceptor cells (Rs) with R1-6 located in the periphery and spanning the whole depth of the ommatidium. In the center, R7 is situated in the distal part of the ommatidium right above R8. Besides the anatomical location, these cells express different photopigments in a well-defined pattern: R1-6 express Rhodopsin 1 (Rh1), R7 express Rh3 and/or Rh4 and R8 express either Rh5 or Rh6 (Rister et al., 2013). To distinguish the contribution of outer versus inner receptor cells, we compared the behavior of *ninaE*^*5*^ (no Rh1) and *rh5^2^;rh3^1^rh4^1^rh6^1^* flies. *NinaE* flies show no difference in E peak timing in LD 20:4 compared to CantonS, *rh5^2^;rh3^1^rh4^1^rh6^1^* mutants in contrast show a significantly advanced E peak, comparable to flies lacking the whole compound eyes (Fig. 2A and 2B). These data suggest that the inner photoreceptors are necessary to mediate long day adaptation.

**Figure 2.**
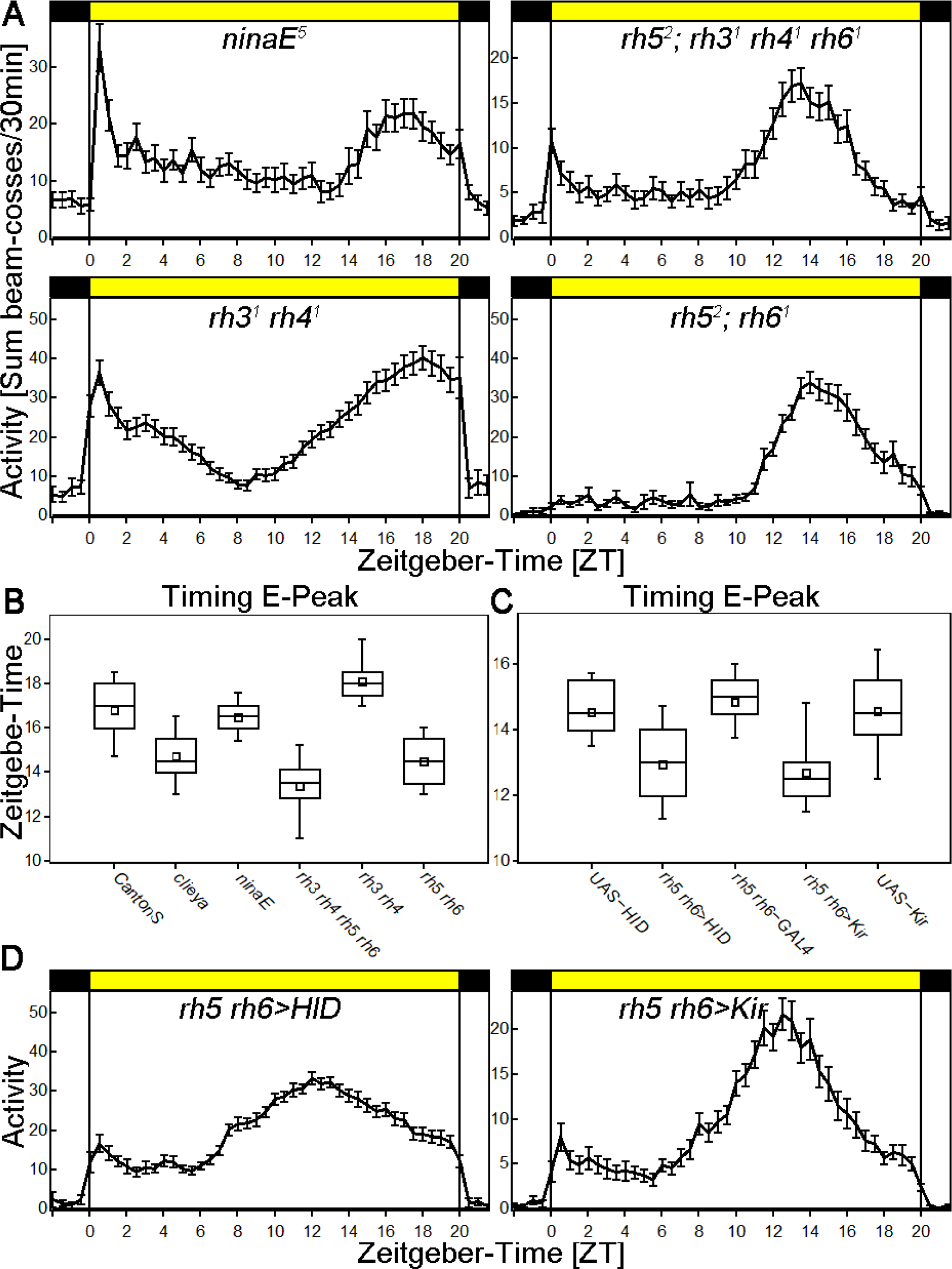
R8 is necessary for long day adaptation. **A** Average activity profiles of photoreceptor mutants recorded in LD 20:4. All genotypes show a bimodal activity pattern with a M peak around lights on and an E peak uncoupled from lights-off. **B** Timing of the E peak in CantonS and all photoreceptor mutants was investigated. One-way ANOVA reveals significant differences between the different genotypes (F_(5,161)_=62.6166, p<0.001). Post-Hoc Tukey comparison shows a significantly advanced E peak in *cli*^*eya*^ mutants compared to CantonS (p=0.001). Similar advances are seen in flies lacking photoreception in both inner receptor cells (*rh5^2^;rh3^1^rh4^1^rh6^1^*, p=0.001) and flies lacking photoreception only in R8 (*rh5^2^;rh6^1^*, p=0.001). There was no difference between *cli*^*eya*^ and *rh5^2^;rh6^1^* mutants (p=0.899), suggesting a prominent role of R8 in long day adaptation. Flies lacking photoreception in R1-6 (*ninaE*^*5*^) show no difference to CantonS (p=0.776), whereas there is a slight but significant delay in flies lacking photoreception in R7 (*rh3^1^rh4^1^*, p=0.001). **C** Timing of the E peak in flies with silenced or ablated R8. One-way ANOVA reveals significant differences between the different genotypes (F_(4,150)_=25.45, p<0.001). Post-Hoc Tukey comparison shows a significantly advanced E peak in flies with silenced R8 (p=0.001 for both controls) and flies with ablated R8 (p=0.001 for both controls). There were neither significant differences between the controls (p>0.775) nor between the two experimental lines (p=0.8926) demonstrating an important role of R8 for long day adaptation. **D** Average activity profiles of of flies with ablated (left) or silenced (right) R8 recorded in LD 20:4. All genotypes show a bimodal activity pattern with a M peak around lights on and an E peak uncoupled from lights-off.

To narrow down the phenotype to a specific inner receptor cell, we monitored the behavior of *rh3^1^rh4^1^* mutants eliminating R7 function and *rh5^2^;rh6^1^* mutants eliminating R8 function. *rh3^1^rh4^1^* mutants behave similar to WT, whereas *rh5^2^;rh6^1^* mutants show an advanced E peak in LD 20:4 indistinguishable from *rh5^2^;rh3^1^rh4^1^rh6^1^* quadruple mutants (Fig. 2A and 2B). This suggests that the rhodopsins of R8 are necessary for WT E peak timing under long photoperiods.

To further confirm the importance of R8 we combined *rh5-GAL4* and *rh6-GAL4* and either silenced these two cell types using UAS-*Kir2.1* or ablated the cells using UAS-*HID*. Immunohistochemistry shows strong specificity of the combined GAL4 lines and a successful ablation of R8 without affecting other photoreceptors in the HID experiment (Suppl. Fig. 1). As with the rhodopsin mutants, ablating or silencing R8 caused a 1.5 hour advance in E activity, confirming an important role for this photoreceptor cell. We further silenced or ablated R7 using a combination of *rh3-GAL4* and *rh4-GAL4*. As expected, there is no effect on E peak timing, confirming the rhodopsin mutant approach data (Suppl. Fig. 2).

As an early E peak might represent a “fast” clock, we monitored the behavior of all mutants in constant darkness (Table 1). We found no correlation between E peak timing and period length, suggesting the long photoperiod phenotype is a true entrainment phenomenon.

**Table 1.** Free-running behavior of WT flies and flies with manipulated visual system. All flies show a period close to 24h.

| | % rhythmic flies | period ± SEM | power ± SEM |
| --- | --- | --- | --- |
| *CantonS* | 84 | 24.3 ± 0.1 | 26.7 ± 1.9 |
| *cli<sup>eya</sup>* | 100 | 23.6 ± 0.1 | 25.4 ± 1.9 |
| *ninaE* | 100 | 23.9 ± 0.1 | 44.2 ± 1.6 |
| *rh5<sup>2</sup>;rh3<sup>1</sup>rh4<sup>1</sup>rh6<sup>1</sup>* | 23 | 23.6 ± 0.1 | 24.0 ± 3.1 |
| *rh3<sup>1</sup>rh4<sup>1</sup>* | 83 | 23.5 ± 0.1 | 25.2 ± 1.2 |
| *rh5<sup>2</sup>;rh6<sup>1</sup>* | 100 | 23.3 ± 0.1 | 23.0 ± 1.8 |
| *UAS-HID* | 100 | 23.9 ± 0.1 | 50.7 ± 2.6 |
| *rh5 rh6>HID* | 100 | 23.4 ± 0.1 | 39.1 ± 3.5 |
| *rh5 rh6-GAL4* | 100 | 23.3 ± 0.1 | 47.6 ± 1.8 |
| *rh5 rh6>Kir* | 100 | 23.3 ± 0.1 | 39.6 ± 1.9 |
| *UAS-Kir* | 100 | 23.6 ± 0.1 | 28.1 ± 2.0 |

### PDF in lLNvs is necessary and sufficient for proper E peak timing

The terminals of R8 directly innervate the medulla, the visual center of the fly, where they sit in close proximity to lLNv arborizations (Schlichting et al., 2016). GRASP experiments between R8 and PDF positive neurons did not give a signal in the medulla, suggesting no direct interaction between the compound eyes and the clock (data not shown). However, electrophysiological data suggest that the visual system activates the PDF expressing ventro-lateral neurons (among others) upon light stimulation (Li et al., 2018; Muraro and Ceriani, 2015). To address the importance of the PDF neurons for long day entrainment, we silenced these cells using *UAS-Kir2.1* (Fig. 3A). Silencing the PDF neurons significantly advances the E peak timing by approximately 1.5h, recapitulating the *cli*^*eya*^ phenotype (Fig. 3B).

**Figure 3.**
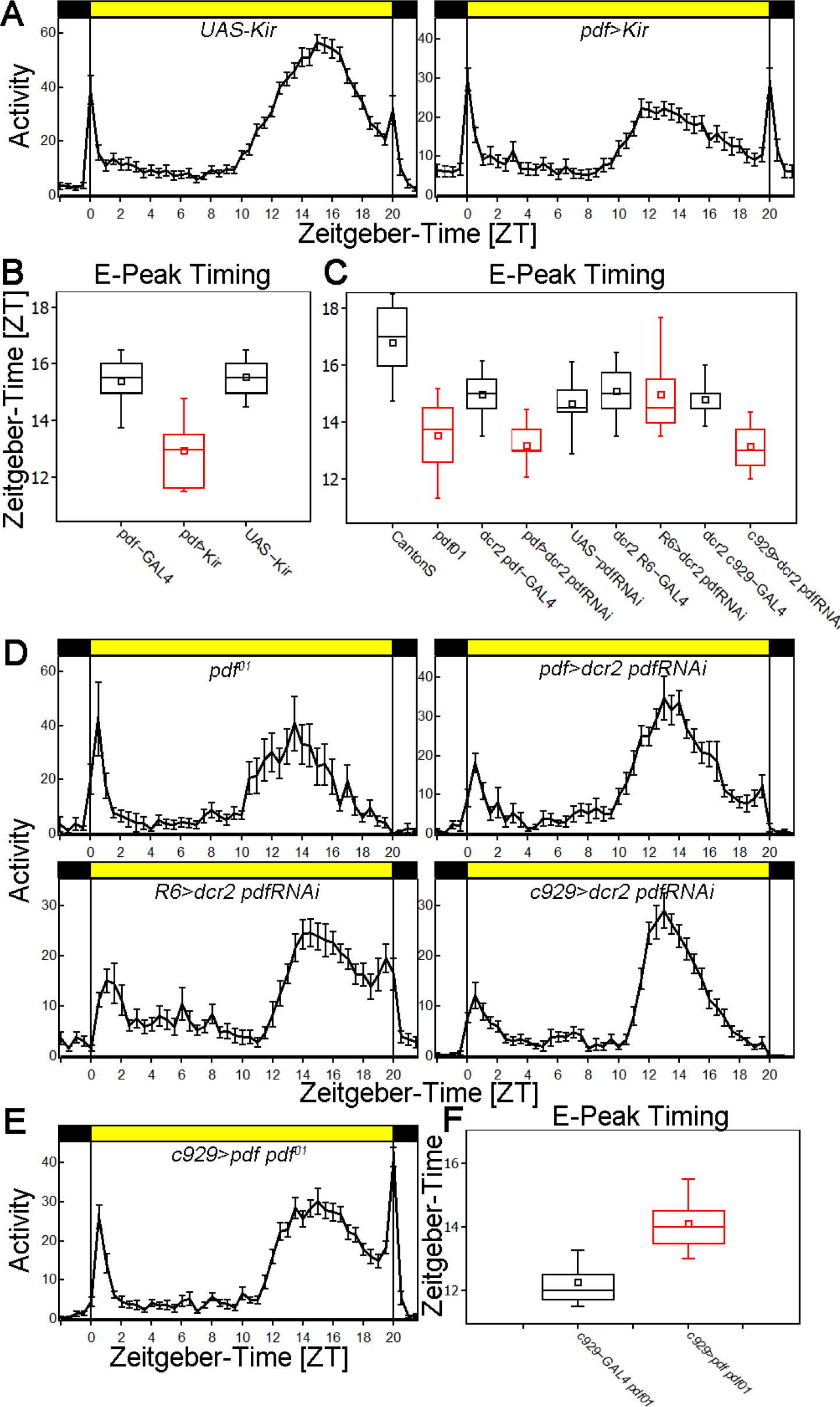
PDF release from the lLNvs is essential for long day adaptation **A** Average activity profiles of control flies (*UAS-Kir*, left) and flies with silenced PDF neurons (*pdf>Kir*, right) in LD 20:4. Flies show a bimodal activity pattern. The timing of the E peak is significantly advanced in *pdf>Kir* flies. **B** Timing of the E peak in flies with silenced PDF neurons including controls in LD 20:4. One-way ANOVA reveals a significant difference between the genotypes (F_(2,89)_=70.1449, p<0.001). Post-hoc Tukey test shows a significantly advanced E peak in *pdf>Kir* compared to both controls (p=0.001 for both). There was no significant difference within control groups (p=0.758). **C** Timing of the E peak in flies with altered PDF expression including controls in LD 20:4. One-way ANOVA reveals a significant difference between the genotypes (F_(8,178)_=28.607, p<0.001). Post-hoc Tukey analysis show that the timing of the E peak is significantly advanced in *pdf^01^* compared to CantonS flies (p=0.001). Similarly, knockdown of PDF using RNAi in both, the sLNvs and lLNvs significantly advanced E peak timing (p=0.001 for both controls). Knockdown of PDF in the sLNvs using *R6-GAL4* had no effect on E peak timing (p=0.899 for both controls), whereas the knockdown in lLNvs using *c929-GAL4* significantly advanced E peak timing (p=0.001 for both controls). There is no difference between *pdf-GAL4* and *c929-GAL4* mediated knockdown (p=0.899) indicating PDF from the lLNvs is necessary for WT behavior. **D** Average activity profiles of *pdf^01^* (top left), PDF-knockdown in all lateral neurons (top right), PDF knockdown in sLNvs (bottom left) and PDF knockdown in lLNvs (bottom right) in LD 20:4. Flies show a bimodal activity pattern. The timing of the E peak is significantly advanced in flies lacking PDF at least in the lLNvs. **E** Average activity profile of PDF rescue in lLNvs in LD 20:4. Rescue of PDF only in lLNvs restores WT behavior. **F** E peak timing of lLNv specific PDF rescue and control. Student’s t-test shows a significant delay of E peak timing (p<0.001) when PDF is rescued in the lLNvs.

PDF is the major neuropeptide of the *Drosophila* clock and is essential to synchronize the different clock neuron clusters with each other (Helfrich-Förster et al., 2007). Previous work showed that PDF from lLNvs is necessary to adapt fly behavior to LD 16:8 (Schlichting et al., 2016). We asked, whether PDF from these neurons is also necessary for proper E Peak timing under even longer days (LD 20:4) and investigated the behavior of *pdf^01^* flies (Yoshii et al., 2009) (Fig. 3B and 3C). As with the *pdf>Kir* experiment, *pdf^01^* flies show an advanced E peak, indicating that PDF signalling to its downstream target neurons is necessary for the delay of the E peak in long photoperiods. To determine, which group of PDF neurons is essential for this behavior, we knocked down PDF using RNAi. PDF knockdown in all PDF-positive cells (sLNvs and lLNvs) results as expected in an advanced E peak compared to both controls (Fig. 3B and 3C). Knockdown only in sLNv using *R6-GAL4* does not advance the timing of the E peak. In contrast, knockdown in lLNvs using *c929-GAL4* completely reproduces universal PDF knockdown (Fig. 3B and 3C) indicating that PDF from the lLNvs is necessary for proper E peak timing under long day conditions.

To address if lLNv-derived PDF is also sufficient for WT behavior, we expressed PDF in the lLNvs using *c929-GAL4* in the *pdf^01^* null mutant background (Fig. 3E). The timing of the E peak was delayed by approximately 1.5h (Fig. 3F), which recapitulates the WT phenotype. The two approaches taken together indicate that PDF from the lLNvs is necessary and sufficient for WT behavior under long photoperiod conditions (Menegazzi et al., 2017).

### E cells show extensive arborizations in the accessory medulla

Previous experiments implicate the 5^th^ sLNv and three CRY-positive neurons in driving/timing the E peak, the evening bout of activity (Grima et al., 2004; Stoleru et al., 2004). These neurons broadly innervate the dorsal part of the brain where they receive glutamatergic input (Guo et al., 2016). They also send fibers into the area of the accessory medulla of the fly brain, the location of the PDF cell bodies and an important pacemaker center in many other insect species (Helfrich-Förster et al., 2007; Schubert et al., 2018).

To determine, whether lLNv-derived PDF could communicate with the E cells in that area, we expressed synaptic markers in the E cells. This was done using a recently identified split-GAL4 line, which expresses only in the three CRY+ LNds and the 5th sLNv (Guo et al., 2017). Whole brain imaging reproduces the previously published projection pattern, showing strong synaptic marker staining in the dorsal brain (Fig. 4A-4D). In the accessory medulla however, we found only weak staining of dendritic and axonal markers. To further illuminate the nature of these E cell fibers, we employed expansion microscopy and focused on this area. The much better resolution indicates that the accessory medulla is indeed densely innervated by E cell fibers, both synaptic as well as dendritic markers (Fig. 4E-4H). It is therefore a likely output as well input region of E cells. The accessory medulla more generally seems to serve as a region of communication between clock neurons, e.g., the lLNvs probably communicate there with the sLNvs via PDF (Choi et al., 2012).

**Figure 4.**
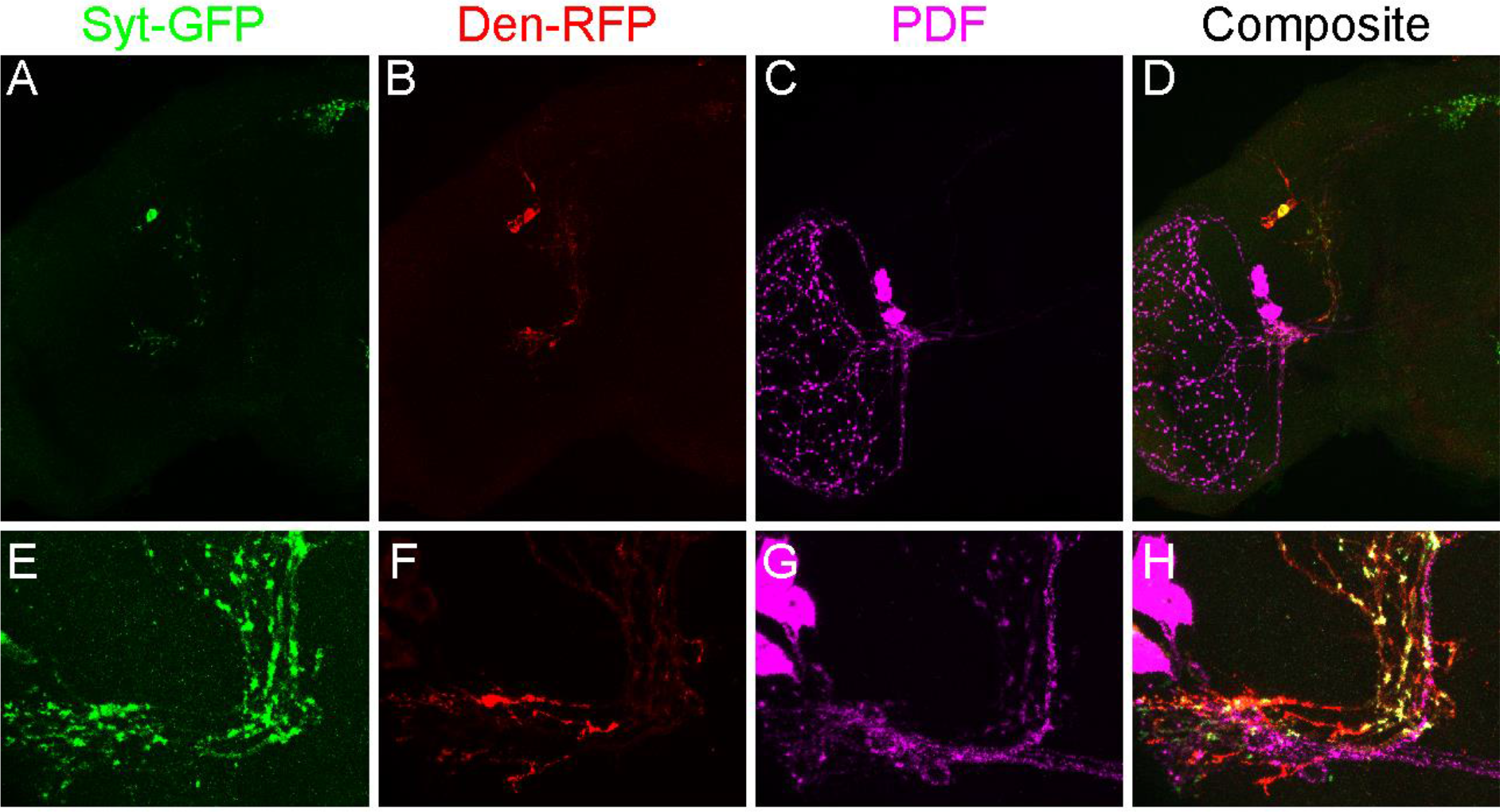
Anatomy of E-cell arborizations in the brain of *Drosophila*. **A-D** Maximum projection of *Spl-E-GAL4>Syt-GFP Den-RFP* flies using regular IHC. **A** Synaptic markers (green) are predominantly present in the dorsal part of the brain. **B** Dendritic markers (red) are more prominent in the accessory medulla. **C** PDF staining (magenta) labels lLNv and sLNv neurons and their projections. **D** Composite of all channels. **E-G** Projections of *Spl-E-GAL4>Syt-GFP Den-RFP* flies in the accessory medulla (aMe) using expansion microscopy. The aMe is densely innervated by E-cells, which express synaptic (green, **E**) and dendritic (red, **F**) markers in close vicinity of PDF neuron fibers (magenta, **G)**. **H** Composite of all channels showing axonal-dendritic nature of E cell arborizations in the aMe.

### PDFR in E cells is necessary and sufficient for proper E peak timing

Loss of PDF or its receptor PDFR (*han*^*5304*^ mutant) causes prominent effects on LD 12:12 and DD behavior: mutant flies show a reduced M anticipation and an early E peak in LD as well as elevated arrhythmicity and a short period phenotype in DD (Hyun et al., 2005; Mertens et al., 2005; Renn et al., 1999). To address the importance of PDFR+ E cells for the long day phenotype, we employed a cell-specific CRISPR/Cas9 strategy with the GAL4/UAS system. We generated an *UAS-PDFRg* line, expressing three independent guides, each targeting the CDS of the *pdfr* gene. To verify the efficiency of this strategy, we expressed *PDFRg* and *Cas9* in most of the clock neuron network using *clk856-GAL4*. It reproduced the *han*^*5304*^ mutant phenotype: flies show a low M anticipation index and an early E peak in LD12:12, and only 37% of the flies are rhythmic with a short period of 22.7 hours in DD (Suppl. Fig. 3).

We then applied the same strategy to long photoperiods. Knocking out PDFR in most clock cells using *clk856-GAL4* reproduced the early E peak phenotype seen in *eyes absent* and *pdf*^*01*^ flies, further indicating that PDF signaling within the clock network is essential for long day adaptation (Fig. 5A and 5B). Knocking out PDFR in E cells using *Mai179-GAL4* reproduced the same behavioral phenotype, i.e., an early E peak under long day conditions (Fig. 5A and 5B). Remarkably, DD behavior was unaffected: 73% of flies were rhythmic with a period of 23.8 ± 0.2h, indicating that E cell PDFR is only required in LD conditions (Suppl. Fig. 4).

**Figure 5.**
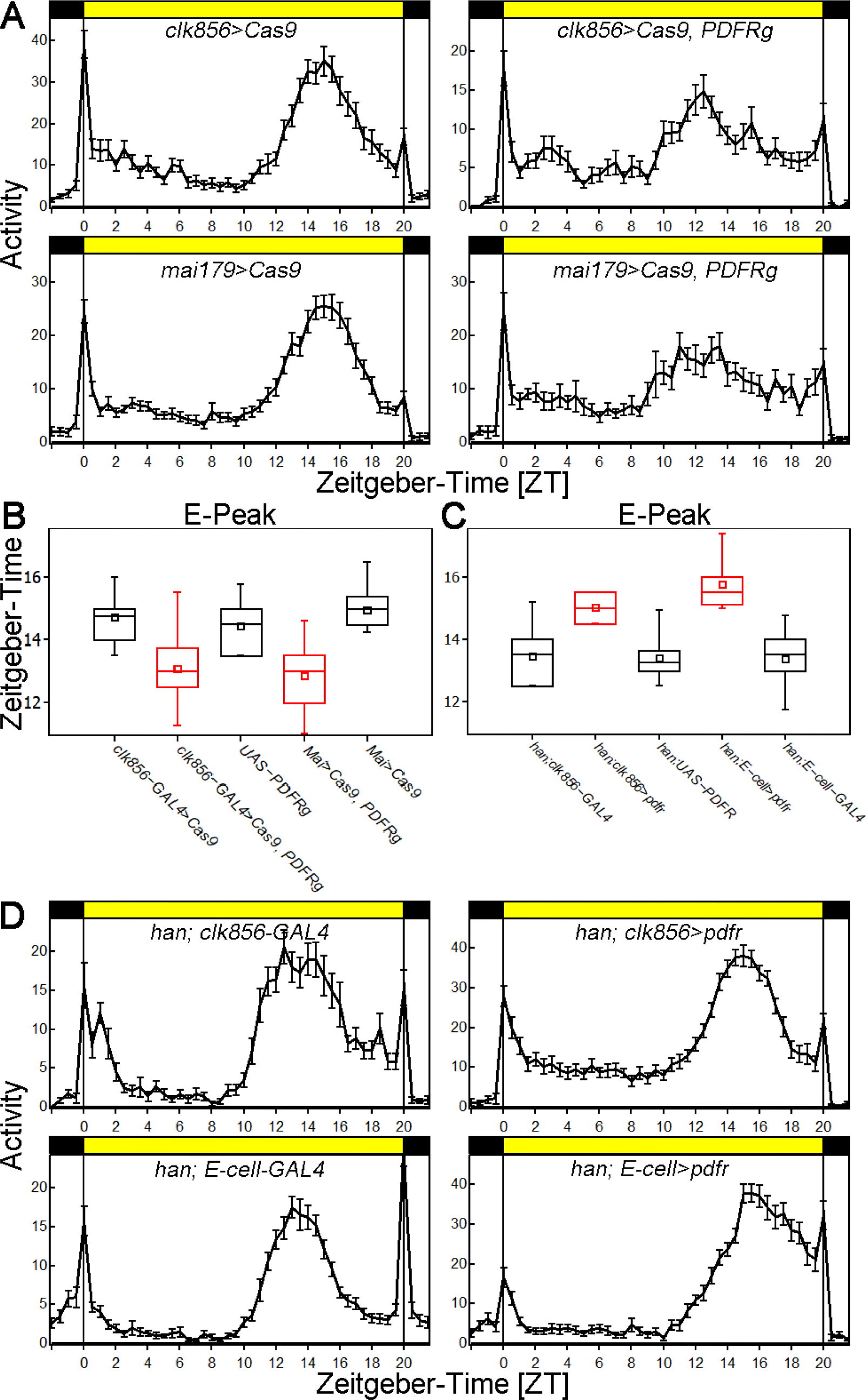
PDFR in E cells is necessary and sufficient for proper E peak timing in LD 20:4. **A** Average activity profiles of *clk856>Cas9* control (top left), PDFR knockout in all clock neurons (top right), *mai179-GAL4*>Cas9 control (bottom left) and PDFR knockout in E-cells using *mai179-GAL4* (bottom right) in LD 20:4. Flies show a bimodal activity pattern with a M peak around lights-on and an E-peak uncoupled from lights-off. **B** Timing of the E peak in PDFR knockout flies including controls in LD 20:4. One-way ANOVA reveals a significant difference between the genotypes (F_(4,131)_=23.945, p<0.001). Post-hoc Tukey test shows a significantly advanced E peak in PDFR knockout in all clock cells compared to both controls (p=0.001 for both). Similarly, knockout of PDFR using *mai179-GAL4* advanced the E peak timing compared to both controls (p=0.001 for both). There was no significant difference between the two knockout strains (p=0.899) showing that PDFR in E cells is necessary for proper E peak timing. **C** Timing of the E peak in PDFR rescue flies including controls in LD 20:4. One-way ANOVA reveals a significant difference between the genotypes (F_(4,125)_=43.358, p<0.001). Post-hoc Tukey test shows a significantly delayed E peak in PDFR rescue in all clock cells compared to both controls (p=0.001 for both). Similarly, rescue of PDFR using *E-cell-Spl-GAL4* delayed the E peak timing compared to both controls (p=0.001 for both). **D** Average activity profiles of *han*^*5304*^;*clk856-GAL4* control (top left), PDFR rescue in all clock neurons (top right), *han^5304^;E-cell-Spl-GAL4* control (bottom left) and PDFR rescue in E-cells using *E-cell-Spl-GAL4* (bottom right) in LD 20:4. Flies show a bimodal activity pattern with a M peak around lights-on and an E-peak uncoupled from lights-off.

Due to a lack of specific driver lines, a previous study implicated E cell PDFR in long day entrainment based on an overlap analysis of much broader driver lines or driver lines with ectopic expression (Schlichting et al., 2016). In contrast, GAL4 lines used here can directly assign E cells to this function. To this end, we rescued PDFR in most of the clock neuron network which delayed the timing of the E peak to WT levels (Fig. 5C and 5D). Rescue of PDFR only in the three CRY+ LNds and the 5^th^ sLNv also delayed the E peak compared to both controls, showing that PDFR in the E cells is indeed sufficient to rescue the E peak timing under long days.

## Discussion

The circadian clock is able to entrain to the changes of day and night, with light being the most important zeitgeber. The adaptation to summer-like days is especially important for insects, as they are prone to predator visibility and even more importantly desiccation. Therefore, the circadian clock has to be plastic and be able to adjust behavior to changing environments. Here we show that *Drosophila* adjusts its behavior to long photoperiods, by delaying its E peak as reported previously (Rieger et al., 2003). This delay allows the animal to reduce its activity during the unfavorable midday, when temperatures are highest. Most interestingly, this phenotype is easily visible even without temperature changes, underscoring the importance of light as the major entrainment cue.

A central finding is that flies lacking the compound eyes show an entrainment deficit, i.e., they have an advanced E peak under long day conditions. Using rhodopsin mutants and by manipulating specific photoreceptors using the GAL4/UAS-system, only R8 is essential for summer day adaptation. Interestingly, R8 was already implicated in the adaptation to nature-like light conditions (Schlichting et al., 2014, 2015).

lLNv arbors in the optic lobe are in close proximity to R8 termini, (Schlichting et al., 2016), where they most-likely interact via cholinergic interneurons. This interaction results in a change of neuronal bursting behavior and hence neuropeptide release (Barber et al., 2016; Muraro and Ceriani, 2015). Indeed, we show here that release of PDF from the lLNvs is necessary and sufficient for proper long day adaptation.

These results are surprising given a recently published study on photic entrainment (Li et al., 2018). It shows that the visual system can activate a broad spectrum of lateral and dorsal neurons; they include sLNvs, lLNvs, ITP+ LNds and DN2s among others. Ablation of PDF neurons left the other neurons responsive to visual input, suggesting a parallel model for clock synchronization, i.e., information from the visual system can be directly transferred to independent classes of clock neurons rather than only via PDF. Nonetheless, we show here that PDF signaling from the lLNvs to the LNds is essential for proper long day adaptation, suggesting that direct transfer of light information to other clock neurons serves other light-mediated functions.

PDF stimulates different adenylate-cyclases and increases cAMP, which leads to the stabilization of PER and consequently a longer period or phase delay (Duvall and Taghert, 2012; Li et al., 2014). Therefore, one view is that removing the compound eyes decreases PDF release from the lLNvs and phase-advances the molecular clock in downstream target neurons like the LNds. This newly discovered “visual system to LNd pathway” might also enhance CRY-mediated photoentrainment: CRY was shown to activate neurons upon stimulation (Fogle et al., 2011), similar to the newly identified light activation of clock neuron pathway (Li et al., 2018). Additional activation of the E cells could therefore contribute to the kinetics of TIM degradation, which was recently shown to be important for the phase advance of E activity under long day conditions (Kistenpfennig et al., 2018).

An intriguing inference of this work is that the principal targets of PDF must change with the environmental conditions. Previous work established the sLNvs as essential for DD rhythmicity (Grima et al., 2004; Stoleru et al., 2004), and recent work shows that these neurons are tightly coupled to the dorsal clock neurons in DD: speeding up the PDF neurons forced the DN1s to follow the short period of the sLNvs (Chatterjee et al., 2018). In LD however, this connection is much weaker, and our cell-type specific CRISPR/Cas9 knockout strategy shows that it is the PDF expressing lLNvs communicate with the LNd neurons (Chatterjee et al., 2018). Our data show that the lLNv to LNd connection is important in LD conditions but does not affect DD behavior.

Importantly, our data not only indicate a qualitative shift of PDF targets between DD and LD but also suggest a quantitative shift of dominance depending on photoperiod or the time of light exposure. In DD, the sLNvs are necessary for rhythmic behavior and show robust cycling in PER oscillations, whereas the lLNvs lose PER rhythms as early as the second day of DD. In equinox conditions, both groups may be relevant (Schlichting et al., 2019): PDF from either the sLNvs or lLNvs is sufficient for WT behavior, and only knockdown in both sets of neurons is able to reproduce the *pdf*^*01*^ mutant phenotype (Shafer and Taghert, 2009). In long photoperiods however, PDF from the lLNvs is necessary and sufficient for proper entrainment, whereas the sLNvs do not contribute to E peak timing (Schlichting et al., 2016). Our data therefore point to a profound circuit switch in response to photoperiod, analogous to the neurotransmitter switching that occurs in the mammalian paraventricular nucleus in response to long photoperiods (Meng et al., 2018).

A similar circuit reorganization might also occur in the principal mammalian brain clock neuron location, the suprachiasmatic nucleus (SCN). We know that light information from the visual system is transferred to cells in the ventral part of the SCN, which expresses VIP (Abrahamson and Moore, 2001). VIP functions similarly to *Drosophila* PDF and is not only important for communication between different parts of the SCN (Aton et al., 2005) but also essential for seasonal encoding. This is because VIP knockout mice show no change in peak width as measured by *in vivo* electrophysiological recordings in response to entrainment to different photoperiods (Lucassen et al., 2012). This suggests that VIP is not only involved in relaying light information beyond the ventral SCN but also in the response to light duration as shown here for PDF in *Drosophila*. It will be interesting to see if different VIP-expressing SCN neurons are involved in this response.

## Competing Interests

The authors declare no competing interests.

## Author Contributions

Conceptualization M.S., C.H.F. and M.R.; Methodology M.S., C.H.F. and M.R.; Investigation M.S., P.W., M.D., P.M. and E.D.B.; Visualization M.S., M.D. and P.W.; Writing - Original Draft M.S. and M.R.; Funding Acquisition M.S., C.H.F. and M.R.; Resources C.H.F. and M.R.

## Acknowledgements

We would like to thank Dr. Leslie Griffith and Dr. Katharine Abruzzi for discussions and comments on the manuscript. Also, we are thankful to Dr. Claude Desplan, Dr. Martin Heisenberg and Dr. Dragana Rogulja for providing fly lines and antibodies. Stocks obtained from the Bloomington *Drosophila* Stock Center (NIH P40OD018537) were used in this study. This work was supported by the German Research Foundation (DFG) Collaborative Research Center (SFB1047 Project A2) and the Howard Hughes Medical Institute (HHMI). M.S. was sponsored by a DFG research fellowship (SCHL2135 1/1).

---

**Suppl Figure 1.**
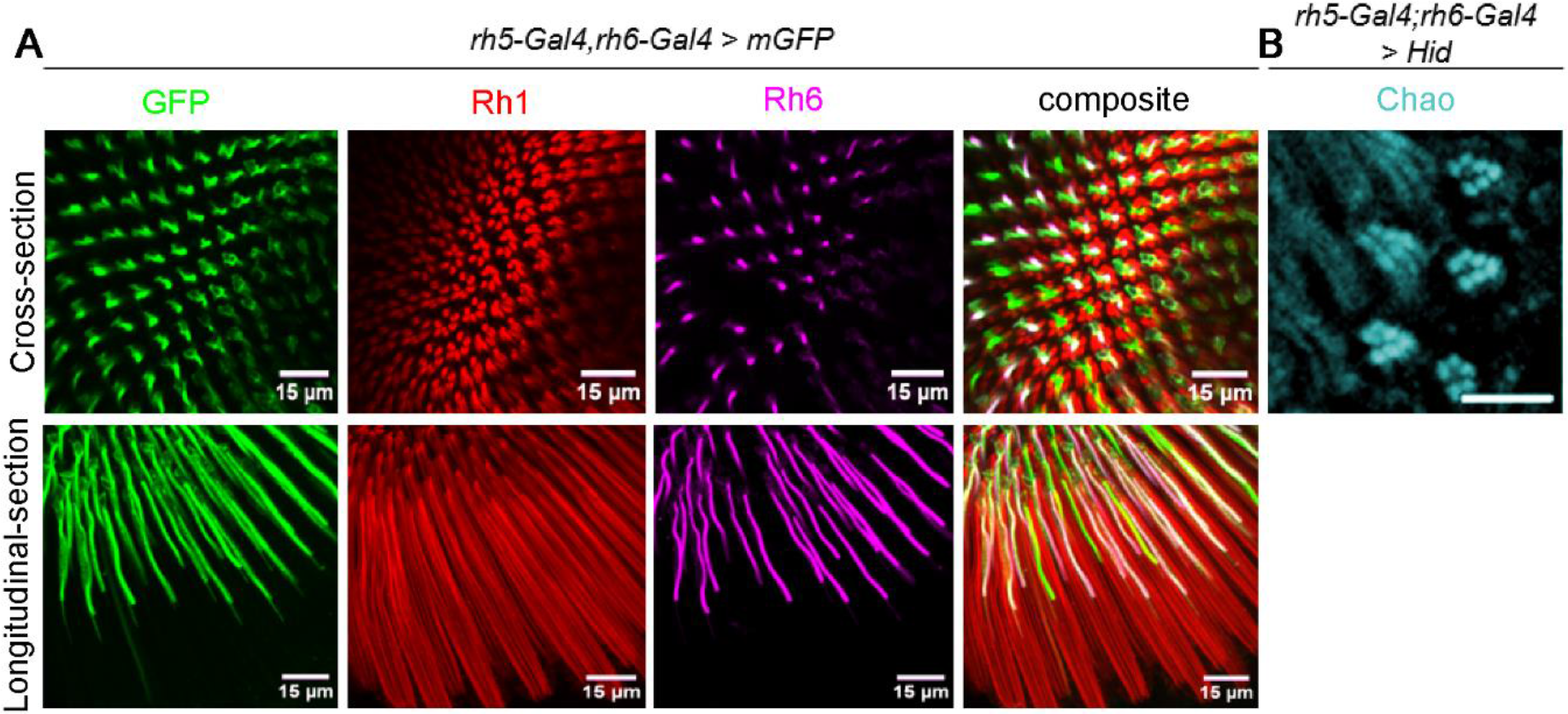
*rh5 rh6-GAL4* combination expresses in all R8s and is able to ablate photoreceptor cells. **A** *rh5rh6>mGFP* stained with anti-GFP (green), anti-Rh1 (red) and anti-Rh6 (magenta). Cross-section through proximal part of the retina shows GFP expression in all inner photoreceptors, but no expression in R1-6. Longitudinal sections show specificity to R8: GFP expression is found in the proximal but not the distal part of the retina. **B** anti-Chaoptin staining of *rh5rh6>HID* (cyan). Chaoptin labels all rhabdomeres. Whereas WT retinas show 7 rhabdomeres per cross section (R1-6 and either R7 or R8), *rh5rh6>HID* flies have only R1-6 remaining, suggesting efficient ablation of R8 in this line. Notably, R1-6 structure seems unaffected.

**Suppl Figure 2.**
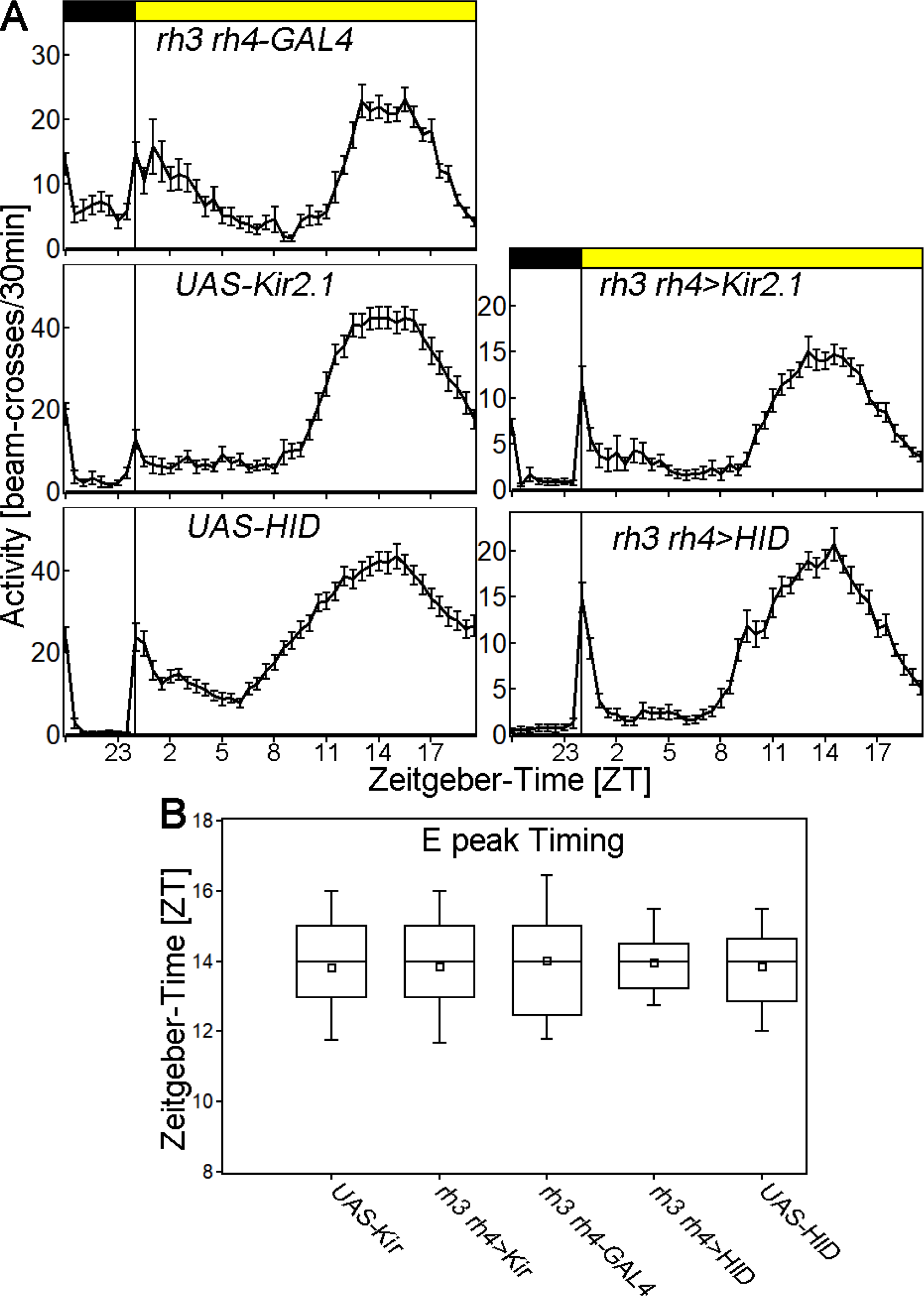
Ablation or silencing R7 has no effect on long day entrainment. **A** Average activity profiles of *rh3rh4-GAL4* control (top left), *UAS-Kir2.1* control (middle panel, left), *UAS-HID* control (lower panel, left) and flies with silenced R7 (middle panel, right) and flies with ablated R7 (lower panel, right) in LD 20:4. Flies show a bimodal activity pattern with a M peak around lights-on and an E-peak uncoupled from lights-off. **B** Timing of the E peak in R7 silenced or R7 ablated flies including controls in LD 20:4. One-way ANOVA with post-hoc Tukey test reveals no significant difference between the genotypes (p>0.144 for pairwise comparisons).

**Suppl Figure 3.**
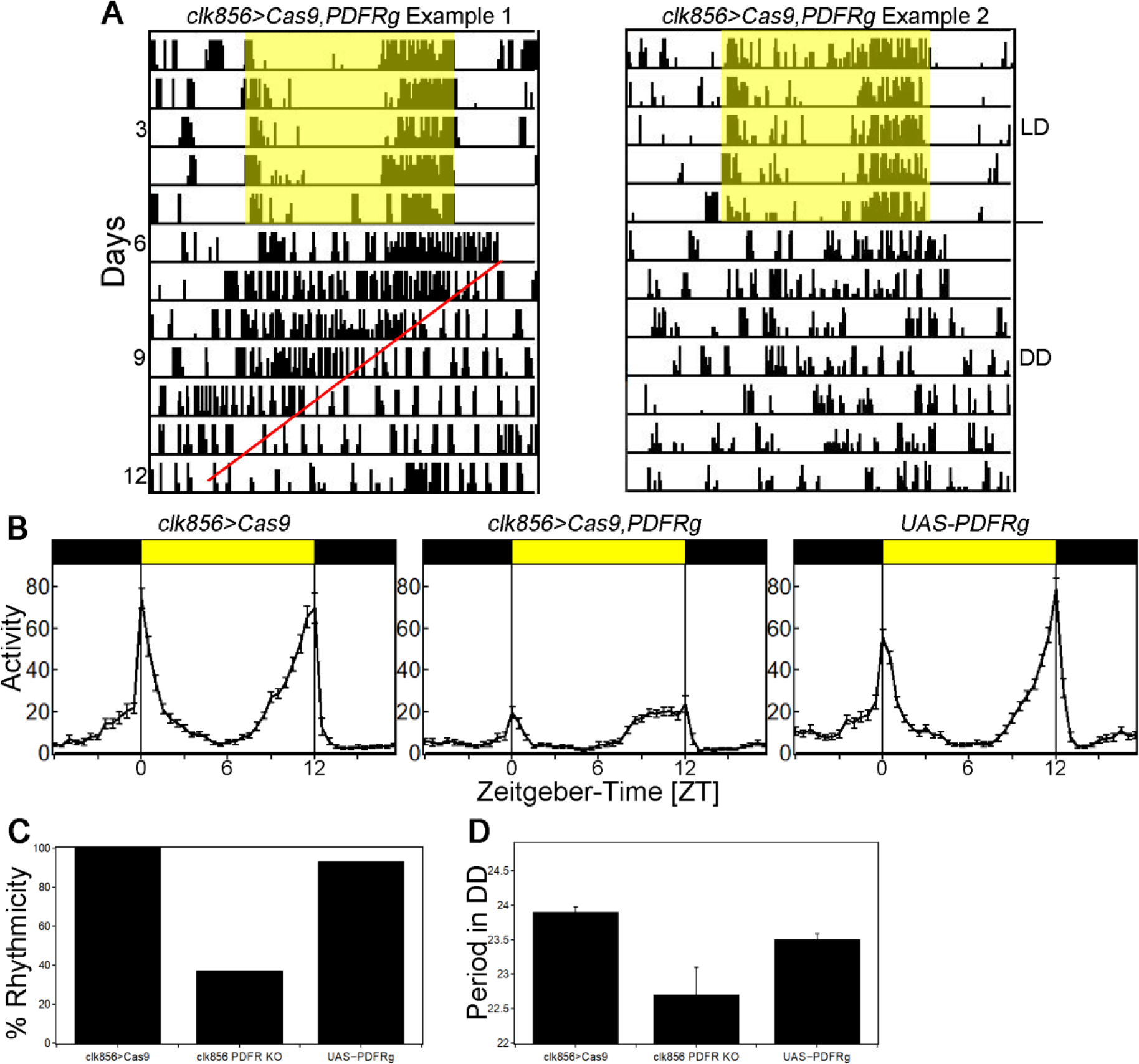
Expression of *PDFRg* and *Cas9* in all clock neurons reproduces *han*^*5304*^ mutant phenotype **A** Example actograms of flies entrained in LD 12:12 for 5 days (indicated by yellow box) and released into constant darkness (DD). Flies either show rhythmic behavior with short period in DD (left example) or arrhythmic behavior (right example). **B** Average activity profiles of *clk856>Cas9* control (left), *clk856>Cas9 PDFRg* (middle) and *UAS-PDFRg* control (right) in LD 12:12. Both controls show a bimodal activity pattern with a M peak around lights-on and an E peak around lights-off. Experimental flies show reduced M anticipation and an advanced E peak comparable to *han*^*5304*^ mutant flies. **C** Percentage rhythmic flies in DD. Both controls show a high percentage of rhythmicity (>90%), whereas less than 40% of PDFR-KO flies remained rhythmic. **D** Free-running period in DD. The period of PDFR-KO flies is about 1h shorter than both controls.

**Suppl Figure 4.**
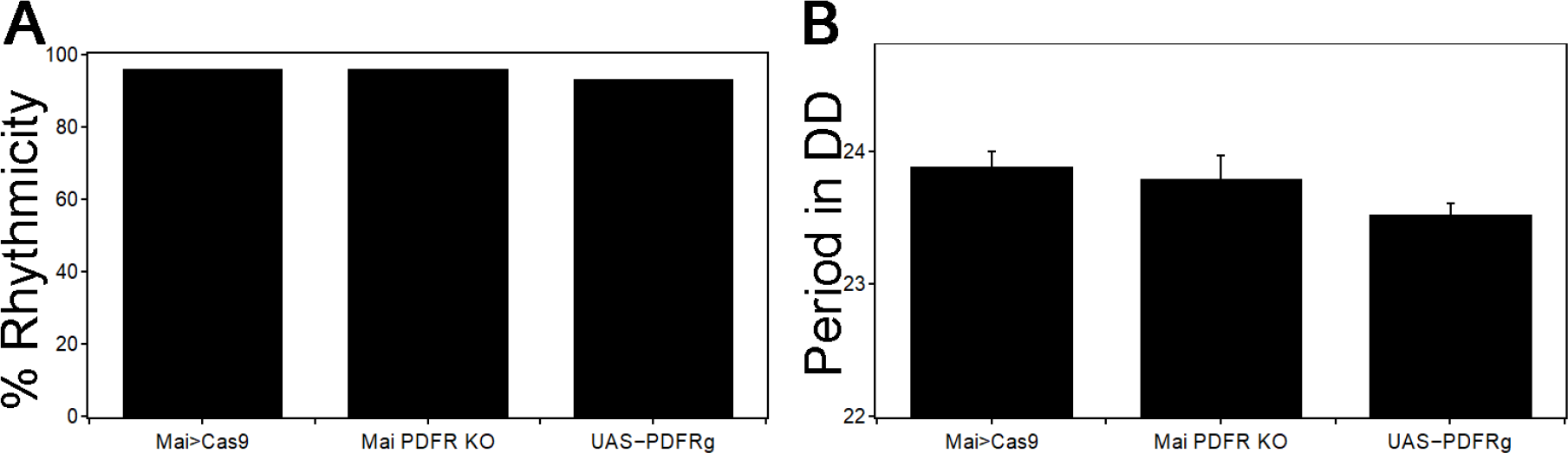
Knockout of PDFR using *mai179-GAL4* has no effect on DD behavior **A** Percentage rhythmic flies in DD. All genotypes show high percentage of rhythmicity (>90%). **B** Free-running period in DD. There is no effect on period length.

